# Verification of extracellular vesicle-mediated functional mRNA delivery via RNA editing

**DOI:** 10.1101/2022.01.25.477620

**Authors:** Masaharu Somiya, Shun’ichi Kuroda

**Affiliations:** SANKEN (The Institute of Scientific and Industrial Research), Osaka University, Ibaraki, Osaka 567-0047, Japan

**Author notes:** To whom correspondence should be addressed., Present Address: Prof. Masaharu Somiya, Ph.D., Department of Biomolecular Science and Reaction, SANKEN (The Institute of Scientific and Industrial Research), Osaka University, Mihogaoka 8-1, Ibaraki, Osaka 567-0047, Japan.

**Keywords:** Alphavirus replicon, CRISPR-Casl3, exosome, extracellular vesicle, mRNA delivery, RNA editing

## Abstract

The secretion and delivery of mRNA by extracellular vesicles (EVs) may contribute to intercellular communications. Several reporter assays have been developed to quantify EV-mediated functional delivery of mRNA into recipient cells. However, mRNA delivery efficiency can often be overestimated by experimental artifacts, resulting in “pseudo-delivery” of reporter proteins rather than mRNA. In this study, we revealed that substantial amounts of reporter proteins expressed in donor cells are secreted into the medium and interfere with the reporter assay. To eliminate this pseudo-delivery, we established a functional RNA delivery assay that employs an RNA editing tool, enabling the verification of *bona fide* delivery of mRNA into recipient cells. The donor cells expressed a reporter gene containing a stop codon in a non-functional open reading frame. After EV-mediated delivery of reporter mRNAs to the recipient cells, guide RNAs and RNA editing enzymes (dCas13b-hADAR2 fusion proteins) correct the RNA sequence and induce the expression of functional reporter proteins in the recipient cells. Using this system, we showed that EVs containing alphavirus-derived replicon successfully delivered functional RNA and expressed the reporter proteins. The RNA delivery assay using RNA editing enables the precise analysis of EV-mediated mRNA delivery.

## INTRODUCTION

Extracellular vesicles (EVs) contain various species of RNAs, such as microRNAs, messenger RNAs (mRNAs), and non-coding RNAs. Extracellular RNAs (exRNA) in EVs are thought to be functionally delivered from donor to recipient cells and regulate biological processes (1, 2). Several studies have demonstrated that EVs deliver mRNAs from donor to recipient cells and functionally translated to the corresponding proteins (1, 3–7). Nevertheless, it is argued that the EV-mediated cargo delivery process might be inefficient (4, 8–10). This controversy is mainly due to the lack of a sensitive and robust bioassay to decipher the delivery mechanism and efficiency of EVs, especially for mRNA-mediated delivery.

Although intercellular shuttling of mRNA is an attractive and plausible mechanism, there are notable caveats in previous studies. For instance, proteins of interest often contaminate the EV preparation, leading to a “pseudo-delivery” of proteins rather than mRNA. Additionally, the transfer of mRNA between the donor and recipient cells is often evaluated by the expression or translation of a reporter gene, such as fluorescence or luminescence proteins, due to its ease of detection and quantification. For the mRNA delivery assay, reporter genes are introduced into the donor cells and expressed, hence, reporter mRNAs are loaded into EVs and secreted. Along with mRNAs, a substantial amount of reporter proteins is expressed in the donor cells and passively loaded into the EVs. Therefore, reporter proteins expressed in the donor cells might be secreted into the conditioned medium due to cell death or other mechanisms. Since reporter proteins are highly sensitive, the contamination of a trace amount of reporter proteins in EV preparations may significantly affect the assay readout. Viral vector preparations are often contaminated with proteins; this contamination leads to a false-positive signal in the target cells, and this process is called “pseudo-transduction” (11, 12). EV-mediated mRNA delivery may be overestimated because of the contamination of reporter proteins in EVs and the conditioned medium. Therefore, there is an urgent need to develop a robust and reliable bioassay to evaluate the intercellular delivery of mRNA from the donor cells to the recipient cells, while excluding the effect of contamination with reporter proteins.

In this study, we developed a reporter gene assay using an RNA editing tool to examine functional RNA delivery. In this assay, upon the EV-mediated delivery of non-functional mRNA into recipient cells, the RNA editing enzyme dCas13b-ADAR2 fusion protein (13) converts the RNA into a functional form, facilitating the detection of mRNA delivery.

## MATERIAL AND METHODS

### Reagents

The NanoLuc substrate, Nano-Glo® Luciferase Assay System, was purchased from Promega Corporation. Synthetic siRNAs were designed and manufactured by Nippon Gene Co., Ltd. and GeneDesign, Inc. The sequences of the antisense strand for siRNA targeting NanoLuc and firefly luciferase were 5’-AUUUUUUCGAUCUGGCCCA-3’ and 5’-UCGAAGUACUCAGCGUAAGTT-3’ (14), respectively.

### Biological Resources

The plasmids used in this study were constructed using a conventional PCR-based method (15). Supplementary Table lists the plasmids used in the present study. Plasmids for VSV-G (Addgene #80054), EGFP (Addgene #89684), and dCas13b-hADDR2 (Addgene #103871) were kindly gifted by Wesley Sundquist, Wilson Wong, and Feng Zhang, respectively.

Human-derived HEK293T cells (RIKEN Cell Bank) were cultured in Dulbecco’s modified Eagle medium (DMEM, high glucose formulation, Nacalai Tesque) containing 10% fetal bovine serum (FBS) and 10 μg/mL penicillin-streptomycin at 37°C in a humidified 5% CO_2_ atmosphere.

### mRNA transfer assay

HEK293T cells were transfected using 25-kDa branched polyethyleneimine (PEI, Sigma) as previously described (8). Briefly, the donor HEK293T cells were seeded in 12 well plates (1–2×10^5^ cells/well, 1 mL/well) or 60 mm dish (1×10^6^ cells/dish, 5 mL/dish) and cultured overnight. The next day, cells were transfected with plasmid DNA (500 ng/well or 2.5 μg/dish, for 12 well plate or 60 mm dish, respectively) and cultured for 2–3 days. After culture, the conditioned medium was collected and centrifuged at 1,500×g for 5 min to remove cell debris. For the isolation of EVs, the conditioned medium was further purified by ultracentrifugation, as previously described (8). Briefly, 1 to 5 mL of supernatant was mixed with PBS and ultracentrifuged (210,000 × g for 70 min by CP100MX ultracentrifuge (Hitachi) and P40ST swing rotor (Hitachi)), then the EV pellet was washed with 12 mL of PBS and centrifuged again, followed by the resuspension in approximately 200 μL of PBS.

Recipient HEK293T cells were transfected one day before the addition of conditioned medium or EVs with corresponding plasmid DNA with or without siRNA, as previously described (8). Recipient HEK293T cells were seeded in 96 well plates ((1–2×10^4^ cells/well, 100 μL/well) and cultured overnight, and then transfected with 100 ng/well of plasmid DNA with or without 1 pmol/well of siRNA. The transfected recipient cells were treated with 100 μL of conditioned medium or 10 μL of purified EVs and cultured for up to 24 h. To evaluate NanoLuc activity, cells were mixed with NanoLuc substrate according to the manufacturer’s instructions. The luminescence signal was quantified using a plate reader (Synergy 2, BioTek).

### Statistical analysis

All experiments were performed in three replicates and conducted at least twice to confirm reproducibility. The data were statistically analyzed using a two-tailed Student’s *t*-test or one-way ANOVA followed by Tukey’s HSD test using the Real Statistics Resource Pack software created by Charles Zaiontz.

## RESULTS

### Reporter proteins in the conditioned medium interfere with the mRNA delivery assay

First, we investigated whether the contamination with reporter proteins in the conditioned medium affected the assay readout. After transfection with plasmids encoding highly bright luciferase NanoLuc originated from *Oplophorus gracilirostris* (16), and the vesicular stomatitis virus glycoprotein (VSV-G), which are known to significantly improve the delivery efficiency of EVs (17, 18), a conditioned medium was collected and added to the recipient human embryonic kidney 293 (HEK293T) cells that were either transfected with siRNAs targeting NanoLuc or control siRNA. If the mRNAs were successfully delivered into recipient cells, strong NanoLuc activity in the recipient cells should be observed, whereas pre-treatment with siRNA targeting NanoLuc should reduce NanoLuc expression in recipient cells by RNA interference mechanism (19). As shown in Figure 1A, we observed a strong NanoLuc activity in recipient cells with VSV-G-conjugated EVs compared to the control (fluorescent protein EGFP). However, we did not observe a significant RNAi-mediated NanoLuc knockdown. These results suggest that rather than NanoLuc mRNAs, the NanoLuc proteins expressed in the donor cells were delivered to the recipient cells, regardless of the presence of VSV-G.

**Figure 1.**
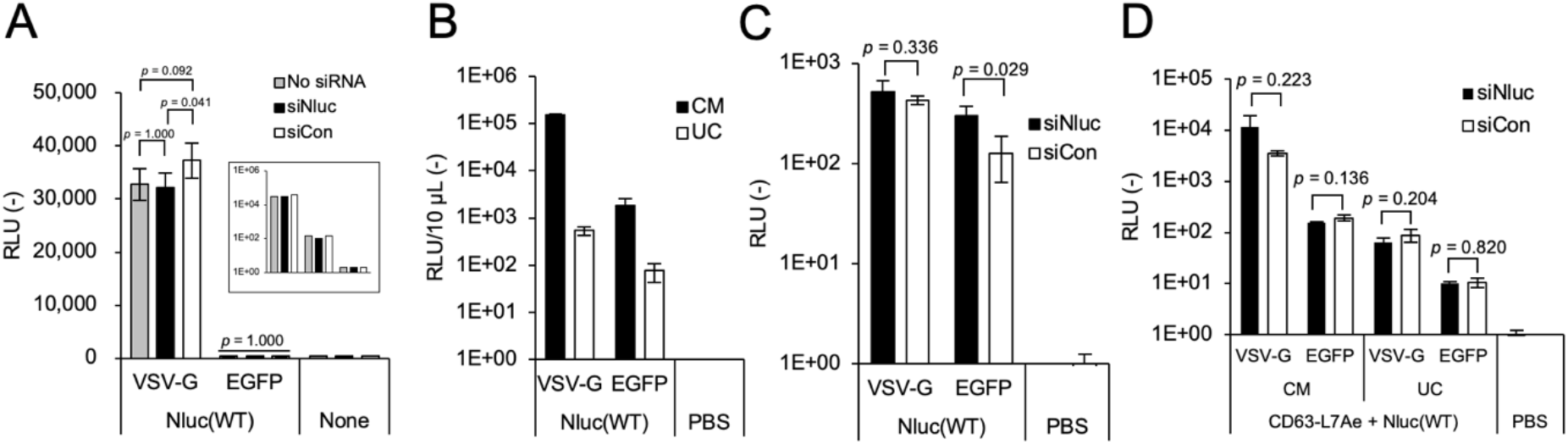
Contamination with reporter NanoLuc proteins in the conditioned medium affected the NanoLuc activity in the recipient cells. (A) NanoLuc activity in recipient HEK293T cells cultured with conditioned medium from donor HEK293T cells. Recipient cells were transfected with or without siRNA targeting NanoLuc (siNluc) or firefly luciferase (siCon, as a negative control) and cultured with 100 μL of conditioned medium for 24 h. Log scale (inset plot) chart was used for the comparison. (B) NanoLuc activity in the conditioned medium (CM) and purified EV preparation after ultracentrifugation (UC). The relative luminescence unit (RLU) per 10 μL of samples is shown. (C) Treatment of recipient HEK293T cells with EVs purified by ultracentrifugation. (D) Conditioned medium (CM) or EVs purified by ultracentrifugation (UC) from donor HEK293T cells expressing RNA loading proteins (CD63-L7Ae), mRNA encoding NanoLuc with tandem C/D box, and VSV-G, were applied to recipient HEK293T cells. N = 3, mean ± SD. The data were analyzed by one-way ANOVA followed by Tukey’s HSD test (A) or Student’s t-test ((C) and (D)).

Next, we isolated EVs from the conditioned medium by ultracentrifugation (Figure 1B). Before ultracentrifugation, the conditioned medium from the donor cells expressing NanoLuc showed high NanoLuc activity, suggesting that a substantial amount of NanoLuc proteins had leaked from the donor cells and contaminated the conditioned medium. Even after ultracentrifugation, the EV preparation showed NanoLuc activity, which was significantly lower than the original conditioned medium, suggesting that despite the removal of the majority of NanoLuc, a substantial amount of NanoLuc proteins remained following ultracentrifugation. Moreover, the addition of purified EVs to the recipient cells significantly increased the NanoLuc activity, which was not affected by the siRNA targeting NanoLuc (Figure 1C). These results show that the reporter proteins contaminating the conditioned medium remained in EV preparations even after purification by ultracentrifugation and that the transporter proteins led to pseudo-delivery in recipient cells.

Previous studies have shown that mRNA can be loaded into EVs and delivered into recipient cells using the EXOTic system which relies on the interactions between the RNA binding proteins L7Ae and specific RNA sequences (kink-turn RNA motif C/D box) (3, 7). We mimicked this system to determine whether this EV-mediated mRNA delivery system was affected by transporter reporter proteins. Donor HEK293T cells were transfected with plasmids encoding CD63-L7Ae, NanoLuc with tandem C/D box at the 3’-UTR, and VSV-G as a delivery enhancer. We then added the transfected donor cell-derived conditioned medium or EVs purified by ultracentrifugation to recipient cells (Figure 1D). Although we observed NanoLuc activity in the recipient cells, this activity was not inhibited by pre-transfection with siRNA targeting NanoLuc, suggesting that the RNA loading system failed to functionally deliver mRNAs into recipient cells, thereby confirming that ultracentrifugation was unable to eliminate the transport of NanoLuc proteins.

### RNA editing for the functional RNA delivery assay

Since contaminations with reporter proteins significantly affect the mRNA delivery assay and lead to an overestimation of the delivery efficiency, we modified the delivery assay, as shown in Figure 2A, to employ a programmable CRISPR-Cas13 system (13). In this assay, we introduced a stop codon at the 12th tryptophan (Trp) operon of the NanoLuc gene. We utilized the RNA editing tool, a fusion protein of catalytically inactive Cas13b (dCas13b) and hADAR2 deaminase domain (E488Q/T375G mutations and lack of C-terminal 984-1090 region), designated as dCas13b-hADDR2, together with the targeting guide RNA (gRNA). The complex of dCas13b-hADDR2 and gRNA edits the amber stop codon (UAG) to UGG, thereby making the mRNAs express the NanoLuc protein. In this system, the donor cells lack the RNA editing mechanism and cannot functionally express NanoLuc; therefore, virtually no transporter NanoLuc protein exists in the conditioned medium and EV preparations. Using this system, we can precisely evaluate functional mRNA delivery while excluding the contamination with transporter reporter proteins from donor cells. This novel mRNA delivery assay was designated as an RNA-editing-based mRNA delivery assay (REMD assay).

**Figure 2.**
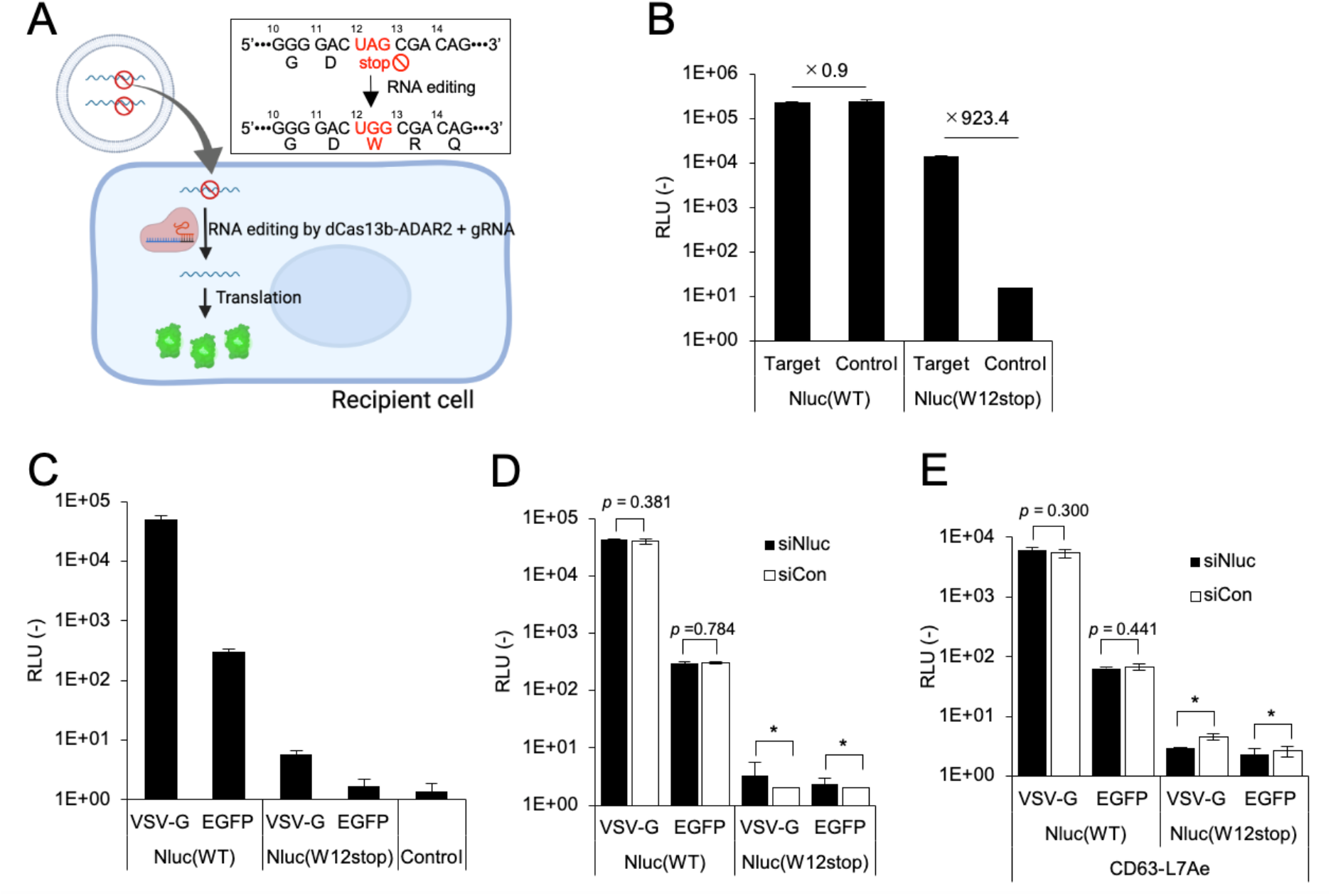
mRNA delivery assay employing CRISPR-Cas13b-based RNA editing tool (REMD assay). (A) Schematic representation of the REMD assay. The upper right inset explains the conversion of the 12th UAG stop codon to the UGG codon of Nluc(W12stop) by dCas13b-hADAR2 and gRNA. (B) Conversion of Nluc(W12stop) mRNA to translationally active mRNA by targeting gRNA in HEK293T cells. Control gRNA targets EGFP(W58stop) and does not affect Nluc(W12stop) mRNA. (C) Validation of REMD assay using donor HEK293T-derived conditioned medium. Recipient HEK293T cells expressing dCas13b-hADAR2 and target gRNA were treated with a conditioned medium for 24 h. (D) REMD assay combined with siRNA treatment. Recipient HEK293T cells transfected with RNA editing tool and siRNA were treated with conditioned medium for 24 h. (E) REMD assay for validation of mRNA delivery using EXOtic system employing NanoLuc(W12stop)-2xC/D box, CD63-L7Ae, and VSV-G. N = 3, mean ± SD. The data were analyzed using Student’s t-test. Asterisks indicate that the statistical analysis was not performed because the luminescence signal was at the background level.

First, we confirmed that the dCas13b-hADDR2 and gRNA complex can precisely edit NanoLuc(W12 stop) mRNA by transfecting HEK293T cells with plasmids encoding NanoLuc(W12 stop), dCas13b-hADDR2, and gRNA, and measuring NanoLuc activity (Figure 2B). The control, NanoLuc without a stop codon, Nluc(WT), was highly expressed regardless of RNA editing, as expected. The luciferase activity of NanoLuc(W12stop) was significantly restored by gRNA targeting Nluc(W12stop), compared to the control gRNA targeting EGFP (W58stop) (20). Thus, we confirmed functional RNA editing using a newly designed gRNA targeting NanoLuc(W12stop).

Next, using the REMD assay, we verified EV-mediated mRNA delivery. Donor cells were transfected with plasmids encoding either Nluc(WT) or Nluc(W12stop), together with VSV-G or EGFP, and the conditioned medium was added to recipient cells expressing RNA editing tools, dCas13b-hADDR2, and gRNA (Figure 2C).

We found that NanoLuc in recipient cells cultured with NanoLuc(W12stop) showed background levels of activity (RLU < 10), whereas recipient cells cultured with NanoLuc(WT) showed high activity (RLU > 10^4^ in the presence of VSV-G). This result indicated that pseudo-delivery of reporter NanoLuc protein in the mRNA delivery assay was successfully excluded using RNA editing tools. We further confirmed the pseudo-delivery of reporter proteins using siRNA (Figure 2D). Knockdown experiments demonstrated that NanoLuc activity in donor cells expressing NanoLuc(WT) was not affected by siNluc, strongly suggesting that transporter NanoLuc proteins significantly affected the mRNA delivery assay.

We further verified the previously reported EV-mediated RNA delivery system (EXOtic device) (3) using the REMD assay. Conditioned medium from donor cells expressing Nluc(W12stop)-2 × C/D box, CD63-L7Ae, and VSV-G was added to the recipient cells, and functional mRNA delivery was evaluated (Figure 2E). Conditioned medium from donor cells expressing NanoLuc (WT) showed significant NanoLuc activity in recipient cells. NanoLuc activity of NanoLuc(WT) samples was derived from transporter reporter proteins, as confirmed by the siRNA targeting NanoLuc. We postulated that the NanoLuc activity observed in recipient cells in a previous study was likely due to contamination with transporter reporter proteins (3). These results indicate that the REMD assay can distinguish *bona fide* mRNA delivery from experimental artifacts owing to the transporter reporter proteins from donor cells.

### Engineered EVs containing alphavirus replicon for functional RNA delivery

We speculated that EV-mediated mRNA delivery has often been overestimated due to contaminations with transporter proteins in the conditioned medium and purified EV preparations. The question remains whether EVs are capable of delivering functional mRNA into recipient cells. Previous studies demonstrated that EVs containing VSV-G and alphavirus replicon RNA can functionally deliver genetic information and express exogenous proteins in recipient cells (21, 22). We used this Venezuelan equine encephalitis virus (VEEV)-derived replicon RNA system (23) encodes the reporter NanoLuc gene under the subgenomic promoter, and supplied VSV-G in *trans* to facilitate the endosomal escape of EVs.

We evaluated the functional mRNA delivery of engineered EVs containing VEEV-derived replicon RNA using the REMD assay (Figure 3A). The VEEV-NanoLuc(WT) samples showed significant NanoLuc activity regardless of the sequence of gRNA, indicating pseudo-delivery of reporter NanoLuc proteins. In contrast, EVs containing VEEV-NanoLuc(W12stop) and VSV-G induced NanoLuc activity in recipient cells expressing the targeting gRNA, suggesting that replicon RNAs were delivered to recipient cells and that RNA editing enzymes converted replicon RNAs into functionally translatable RNAs. The EVs containing VEEV-NanoLuc(W12stop) without VSV-G failed to functionally deliver the replicon RNA, suggesting that endosomal escape and cytoplasmic delivery of RNAs can be successfully achieved by engineering EVs with membrane fusion proteins.

**Figure 3.**
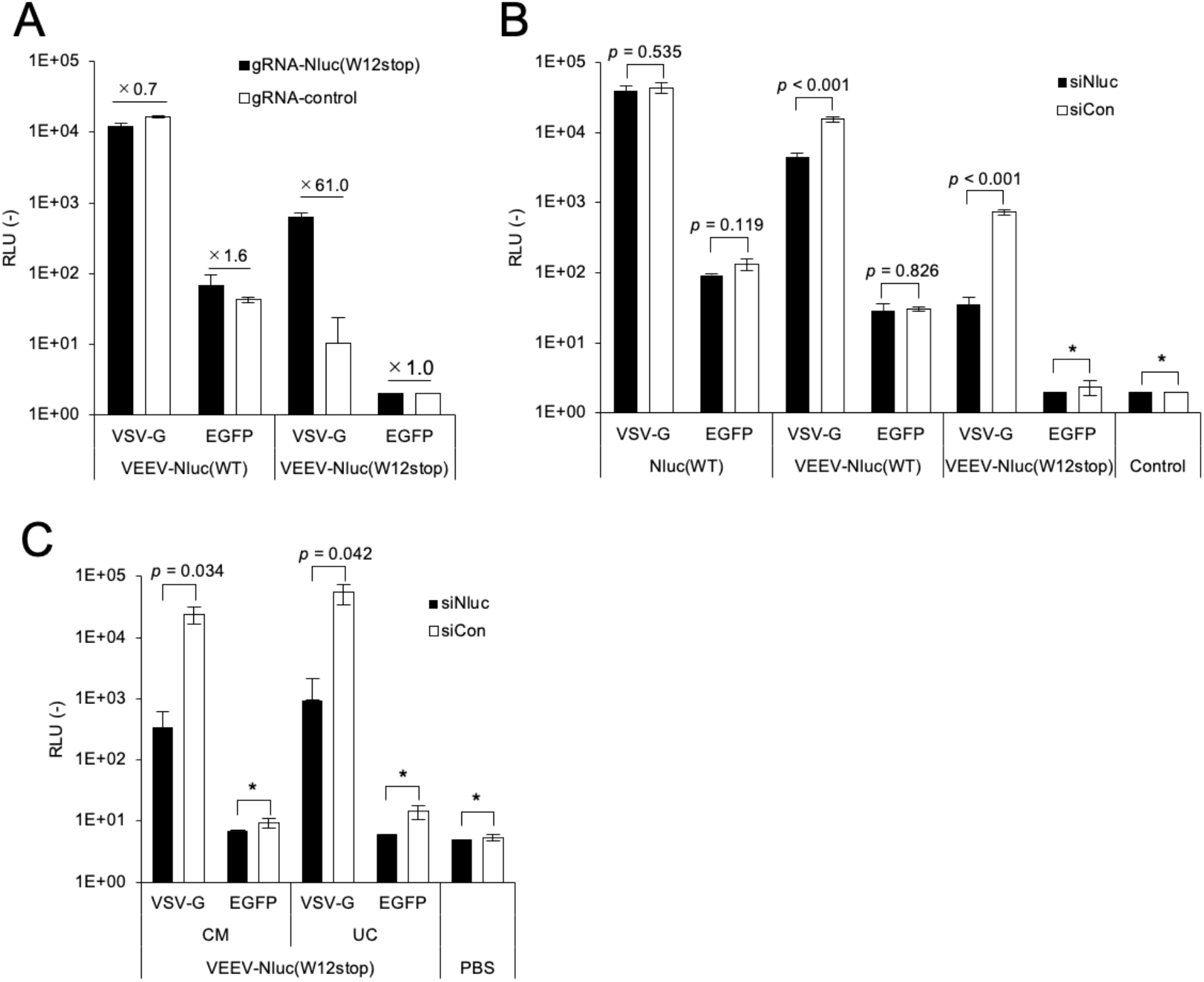
Evaluation of EV-mediated replicon RNA delivery by REMD assay. (A) Donor HEK293T cells were transfected with plasmid encoding VEEV-Nluc(WT) or VEEV-Nluc(W12stop) together with VSV-G or EGFP. The supernatant from the donor cells was added to recipient HEK293T cells expressing dCas13b-hADAR2 and targeting or control gRNA and cultured for 24 h. Numbers above bars represent the fold-increase of RLU by the target gRNA against the control gRNA. (B) Supernatant from transfected donor HEK293T cells was added to the recipient HEK293T cells transfected with plasmid encoding dCas13b-hADAR2 and targeting gRNA, and siRNA targeting NanoLuc (siNluc) or firefly luciferase (siCon). (C) Conditioned medium (CM) or ultracentrifugation-purified EVs (UC) from transfected donor HEK293T cells was added to the recipient HEK293T. Due to the relatively lower luminescence signal in this experiment, the sensitivity setting of the instrument was increased, therefore the values are not comparable to other data. N = 3, mean ± SD. The data were analyzed using Student’s t-test. Asterisks indicate that the statistical analysis was not performed because the luminescence signal was at the background level.

We further verified the engineered EV-mediated functional delivery of replicon RNA using siRNA (Figure 3B). The activity of NanoLuc in recipient cells treated with EVs containing VEEV-NanoLuc(WT) and VSV-G was reduced by 70% (*p* < 0.001), suggesting that a fraction of this activity was due to translation of functional RNAs; however, transporter proteins still contributed to NanoLuc activity in recipient cells. In contrast, over 95% of NanoLuc activity by the EVs containing VEEV-NanoLuc(W12stop) and VSV-G was suppressed by siRNA (*p* < 0.001), strongly suggesting that NanoLuc activity was exclusively driven by the functional delivery of replicon RNA. As well as conditioned medium, purified EVs containing replicon RNA and VSV-G induced the reporter gene expression in the recipient cells, and the reporter gene expression was strongly inhibited by siRNA targeting NanoLuc (Fig. 3C). Collectively, these results confirm that the combination of the REMD assay and knockdowns using siRNA can detect EV-mediated functional RNA delivery into recipient cells.

## DISCUSSION

In this study, we demonstrated that the reporter gene-based assay for EV-mediated mRNA delivery is easily affected by transporter reporter proteins. Therefore, we established a REMD assay that employs an RNA editing tool to exclude the pseudo-delivery of transporter proteins from donor cells. The key feature of the REMD assay is that the mRNA of the reporter gene is translationally inactive within the donor cells, and becomes translationally active in recipient cells through the conversion of a stop codon via CRISPR-Cas13b-mediated RNA editing. A previous study has shown that the codon replacement of the reporter gene is useful for studying RNA editing efficiency using the fluorescence protein EGFP with W58stop mutation (20). Compared to other fluorescence reporters, such as EGFP, NanoLuc shows higher sensitivity and a broader dynamic range. Therefore, we selected the NanoLuc reporter for the sensitive and robust evaluation of EV-mediated mRNA delivery in our REMD assay.

In this study, we showed that HEK293T-derived EVs could not deliver reporter mRNAs into recipient HEK293T cells. The activity of NanoLuc seen in recipient cells was derived from transporter proteins rather than *de novo* proteins translated from mRNA within recipient cells. Even after ultracentrifugation, transporter proteins remained in the EV preparation and significantly affected the assay readouts. As previously described by our group and other research groups, EVs have a low cargo delivery efficiency (4, 8, 9, 24, 25). Especially, Albanese et al. demonstrated that EVs from five different human cell lines could not deliver their cargo against 17 different recipient cell lines unless donor cells express fusogenic proteins (i.e. VSV-G)(4). Thus, we concluded that, in general, EVs hardly deliver mRNA cargo into recipient cells. Conversely, some reports have argued that EVs have the potential to deliver RNA cargo into recipient cells and that mRNA is functionally translated. For example, Kojima et al. demonstrated that mRNA can be packaged into EVs using RNA-protein interactions, and the resultant EVs can functionally deliver the reporter mRNA into recipient cells with the help of production and delivery enhancers (3). However, using the REMD assay, we could not validate the findings of Kojima et al., and we speculated that their assay readout was overestimated due to the transporter reporter proteins in the EV preparation. It should be noted that Kojima et al. used combinations of EV-producing enhancers (STEAP3, NadB, and SDC4) and delivery enhancers (RVG-lamp2b and Cx43-S368A) together with a CD63-L7Ae and NanoLuc-C/D box. In contrast, we simplified the system using VSV-G as a delivery enhancer. The difference in the RNA delivery systems between these studies must be considered when interpreting the results. Furthermore, EV-mediated cargo delivery may be influenced by many factors, including the donor-recipient pair, preparation methods of EVs, and culture conditions of donor cells. Therefore, our findings suggesting that EVs cannot deliver mRNAs should not be generalized in a broad biological context.

We demonstrated that EVs containing viral glycoprotein (VSV-G) and alphavirus replicon successfully delivered RNAs and that a substantial amount of the reporter NanoLuc protein was expressed in recipient cells. The alphavirus replicon RNA replicates within the budding structure, called spherules, at the cell surface (26), and it was assumed that EVs containing both the alphavirus replicon and VSV-G can be released into the extracellular space (21). Upon EV-mediated cytoplasmic delivery, the replicon RNA can be self-amplified in the cytoplasm of recipient cells and strongly express the exogenous gene under the subgenomic promoter. The competency of the self-amplification of the alphavirus replicon makes it an ideal gene delivery vector because a small amount of replicon RNA can highly express the exogenous gene. This suggests that EVs containing an alphavirus replicon are an alternative strategy to successfully deliver functional RNA and express therapeutic genes in target recipient cells. Additionally, the glycoprotein of EVs can be replaced in a process called “pseudotyping” to alter the tropism of the target as conventional viral vectors (21, 27). Thus, pseudotyped EVs containing replicon RNA can target various tissues or cells.

In conclusion, our novel REMD assay is capable to investigate EV-mediated mRNA delivery by excluding artifacts derived from contamination. To date, the lack of a feasible and reliable reporter assay had hampered the understanding of the efficiency and mechanism of EV-mediated mRNA delivery. The REMD assay would help to settle the controversy of whether EV-mediated mRNA delivery practically contributes to intercellular communication and is physiologically relevant. Furthermore, efficient mRNA delivery is one of the key requirements for the development of novel modalities of therapeutics and vaccines based on the mRNA (28, 29). The REMD assay could be expanded to a broader context from fundamental research on EV-mediated mRNA delivery to validation of therapeutic delivery of mRNA.

## DATA AVAILABILITY

All data in this study were described in the main text.

## ACKNOWLEDGMENTS

We extend our gratitude to Yumi Yukawa for providing technical assistance in plasmid construction. All illustrations used in this study were created using BioRender.com.

