## Supplementary Table for "Verification of extracellular vesicle-mediated functional mRNA delivery via RNA editing"

| Number | Name | Insert | Availability | Reference |
|---|---|---|---|---|
| 1 | pCMV-VSV-G-Myc | VSV-G, C-terminal Myc tag | Addgene #80054 | 1 |
| 2 | pCAG-EGFP | EGFP | Addgene #89684 | 2 |
| 3 | pC0055-CMV-dPspCas13b-GS-ADAR2DD(E488Q/T375G)-delta-984-1090 | dPspCas13b-hADAR2DD(E488Q/T375G) | Addgene #103871 | 3 |
| 4 | pcDNA3.1-Nluc | Nluc | Addgene #180501 | This study |
| 5 | pcDNA3.1-Nluc(W12stop) | Nluc(W12stop) | Addgene #180502 | This study |
| 6 | pCMV-CD63-L7Ae | human CD63-L7Ae | Addgene #180503 | This study |
| 7 | pCIneo-Nluc-2xC/D box | Nluc and tandem C/D box at 3' UTR | Addgene #180504 | This study |
| 8 | pCIneo-Nluc(W12stop)-2xC/D box | Nluc(W12stop) and tandem C/D box at 3' UTR | Addgene #180505 | This study |
| 9 | PasCas13b-gRNA-Nluc(W12stop) | guide RNA for PasCas13b, targeting Nluc(W12stop) | Addgene #180506 | This study |
| 10 | PasCas13b-gRNA-EGFP(W58stop) | guide RNA for PasCas13b, targeting EGFP(W58stop) | Addgene #180507 | This study |
| 11 | pcDNA3.1-EGFP(W58stop)-HiBiT | EGFP(W58stop) | Addgene #180508 | This study |
| 12 | pCMV-VEEV-Nluc | VEEV replicon and Nluc | No* | This study |
| 13 | pCMV-VEEV-Nluc(W12stop) | VEEV replicon and Nluc(W12stop) | No* | This study |

1 J. Votteler et al., “Designed proteins induce the formation of nanocage-containing extracellular vesicles.,” Nature, vol. 540, no. 7632, pp. 292–295, Nov. 2016, doi: 10.1038/na
2 B. H. Weinberg et al., “Large-scale design of robust genetic circuits with multiple inputs and outputs for mammalian cells,” Nature Biotechnology, vol. 35, no. 5, pp. 453–462, 2
3 D. B. T. Cox et al., “RNA editing with CRISPR-Cas13,” Science, vol. 358, no. 6366, pp. 1019–1027, Nov. 2017, doi: 10.1126/science.aaq0180.

* Not depositted to Addgene
DNA sequence of plasmid is available upon request
